# Encephalomyocarditis virus protein 2B* antagonises innate immune signalling by interacting with 14-3-3 protein family members

**DOI:** 10.1101/2024.07.31.605952

**Authors:** Samantha K. Nguyen, Stephen Holmes, Henry G. Barrow, Nina Lukhovitskaya, Aminu S. Jahun, Iliana Georgana, Laura G. Caller, James R. Edgar, Edward Emmott, Andrew E. Firth, Hazel Stewart

**Affiliations:** Department of Pathology, University of Cambridge, Cambridge, United Kingdom; Centre for Proteome Research, Department of Biochemistry, Cell & Systems Biology, Institute of Systems, Molecular and Integrative Biology, University of Liverpool, Liverpool, United Kingdom

**Keywords:** Picornavirus, EMCV, interferon beta, cardiovirus, ribosome frameshifting, innate immunity, proteomics, 14-3-3 proteins, 2B*

## Abstract

Encephalomyocarditis virus (EMCV) has for decades served as an important model RNA virus. Although most of the EMCV proteins are obtained via proteolytic cleavage of a long polyprotein, 2B* is expressed from a short overlapping open reading frame via an unusual protein-stimulated temporally dependent ribosomal frameshifting mechanism. The function of 2B* has not yet been characterised, though mutant viruses that are unable to express 2B* have a small plaque phenotype. Here we show that 2B* binds all seven members of the 14-3-3 protein family during virus infection. Binding is dependent on the 2B* C-terminal sequence RRNSS. IFN-β and IL-6 signalling are impeded following overexpression of 2B* but not a truncated version lacking the RRNSS residues, thus suggesting a 14-3-3-dependent role for 2B* in inhibiting MAVS signalling. We also find that this function is distinct from the effect of 2B* on plaque size, as a virus in which 2B* was similarly truncated exhibited near-wildtype plaque size, thus indicating that 2B* also harbours additional functions. This work provides the first identification of a role of 2B* in innate immune antagonism and expands our knowledge of the protein complement of this important model virus.

**IMPORTANCE:** Encephalomyocarditis virus (EMCV) infects a range of species, causing economically important reproductive disorders in pigs and encephalitis and myocarditis in rodents. Due to its wide host range, it is an important model pathogen for investigating virus-host interactions. EMCV expresses an accessory protein, 2B*, from an overlapping open reading frame via an unusual ribosomal frameshifting mechanism. Although the frameshifting mechanism has been established, the function of the 2B* protein had not previously been explored. Here, we determined the host proteins to which 2B* binds and found that it specifically binds to the entire 14-3-3 protein family which, among other roles, contribute to the innate immune response to viral infection in mammalian cells. This interaction requires a specific stretch of amino acids at the end of 2B*. By interacting with the 14-3-3 proteins, 2B* blocks immune response activation. Thus, 2B* is a novel antagonist of innate immunity.

## INTRODUCTION

Encephalomyocarditis virus (EMCV) is a positive-sense single-stranded RNA virus in the genus *Cardiovirus* of the family *Picornaviridae*. EMCV causes encephalitis and myocarditis in a variety of species, as well as reproductive disorders in pigs. The genome is approximately 8 kb in length and encodes a long polyprotein which is cleaved to produce 12 mature virus proteins. Although most of the polyprotein processing is performed by the viral 3C protease, separation between 2A and 2B occurs via the StopGo mechanism whereby a specific amino acid sequence ending in NPGP co-translationally prevents formation of a peptide bond between the G and the final P (1). Just 12 codons downstream of the 2A|2B junction, lies a programmed ribosomal frameshifting (PRF) site at which a proportion of ribosomes are stimulated to make a −1 nt shift into the 117-codon overlapping 2B* open reading frame (ORF) (2). The shift site comprises a G GUU UUU heptanucleotide (spaces separate polyprotein-frame codons), and the stimulator comprises a 3′ RNA structure that binds the viral 2A protein to form an RNA:protein complex that impedes ribosome processivity (3, 4). Ribosomes which make a −1 nt shift can more efficiently remove 2A and unwind this RNA structure to continue translation (3). PRF leads to the production of the 129 amino acid "transframe" protein 2B*, and prevents translation of the downstream nonstructural proteins. The efficiency of PRF is regulated by the increasing concentration of 2A over time, with the percentage of frameshifting ribosomes increasing from 0% at 2 h post infection (h p.i.) to ∼70% at 6–8 h p.i. (3). Thus both the expression level of 2B* and the ratio of structural to nonstructural proteins are temporally controlled.

Although the amino acid sequence of 2B* is highly conserved between EMCV isolates, it is not encoded by other picornaviruses, even the closely related cardiovirus, Theiler’s murine encephalomyelitis virus (TMEV). Although PRF does occur at the same site in TMEV, it leads to the expression of a peptide only 14 residues in length that has no known function (5). Therefore, in TMEV it is thought that PRF is used purely as a "ribosome sink" to temporally control the ratio of structural to nonstructural protein synthesis (5, 6). 2B* therefore represents an entirely uncharacterised protein, present in a subgroup of cardioviruses. The loss of 2B* has been associated with reduced viral plaque size (2, 3), however the molecular mechanisms responsible for this have not yet been elucidated.

Here, we modified the EMCV genome to encode an HA-tagged 2B* protein. Using this virus, we identified the host protein binding partners of 2B* and found that they include the entire family of 14-3-3 proteins. A short linear motif at the C terminus of 2B* is essential for 14-3-3 binding. Interaction of 2B* with the 14-3-3 members reduced transcription of both *IFNB1* and *IL6* in addition to interferon stimulated gene (ISG) transcripts, in accordance with recently published results describing a role of 14-3-3 proteins in promoting antiviral immunity (7–10). This antagonism of the innate immune system, via sequestration of the 14-3-3 family, is distinct from the previously described role of 2B* in promoting plaque size, thus indicating that 2B* possesses multiple functions which contribute to efficient EMCV replication and transmission.

## RESULTS

### **Identification of a 2B*KO EMCV mutation that does not affect viral frameshifting or replication**

As the ORF of 2B* overlaps the coding sequence of 2B, we first sought to confirm that mutations applied to create a 2B* knock-out (2B*KO) virus do not affect either viral PRF efficiency or RNA replication. Several 2B*KO EMCV mutants have been described previously (2, 11), one of which possesses two stop codons immediately after the RNA stem-loop, which are synonymous in the 2B reading frame (WT-PTC mutant of Ling et al., 2017). In this virus, PRF results in expression of a severely truncated, 29 residue N-terminal fragment of the 129 residue 2B*.

As PRF in EMCV directs ribosomes out of the polyprotein reading frame into the 2B* reading frame, any effect of the 2B*KO mutation on PRF efficiency would affect the ratio of structural to nonstructural protein synthesis at late stages of infection, potentially overshadowing any phenotype caused by the loss of 2B*. While previous work with a metabolic labelling assay (11) indicated that the same 2B*KO mutation did not have a measurable effect on frameshifting, we sought to confirm this with an alternative, dual luciferase-based assay (Figure 1A, 1B).

**Figure 1:**
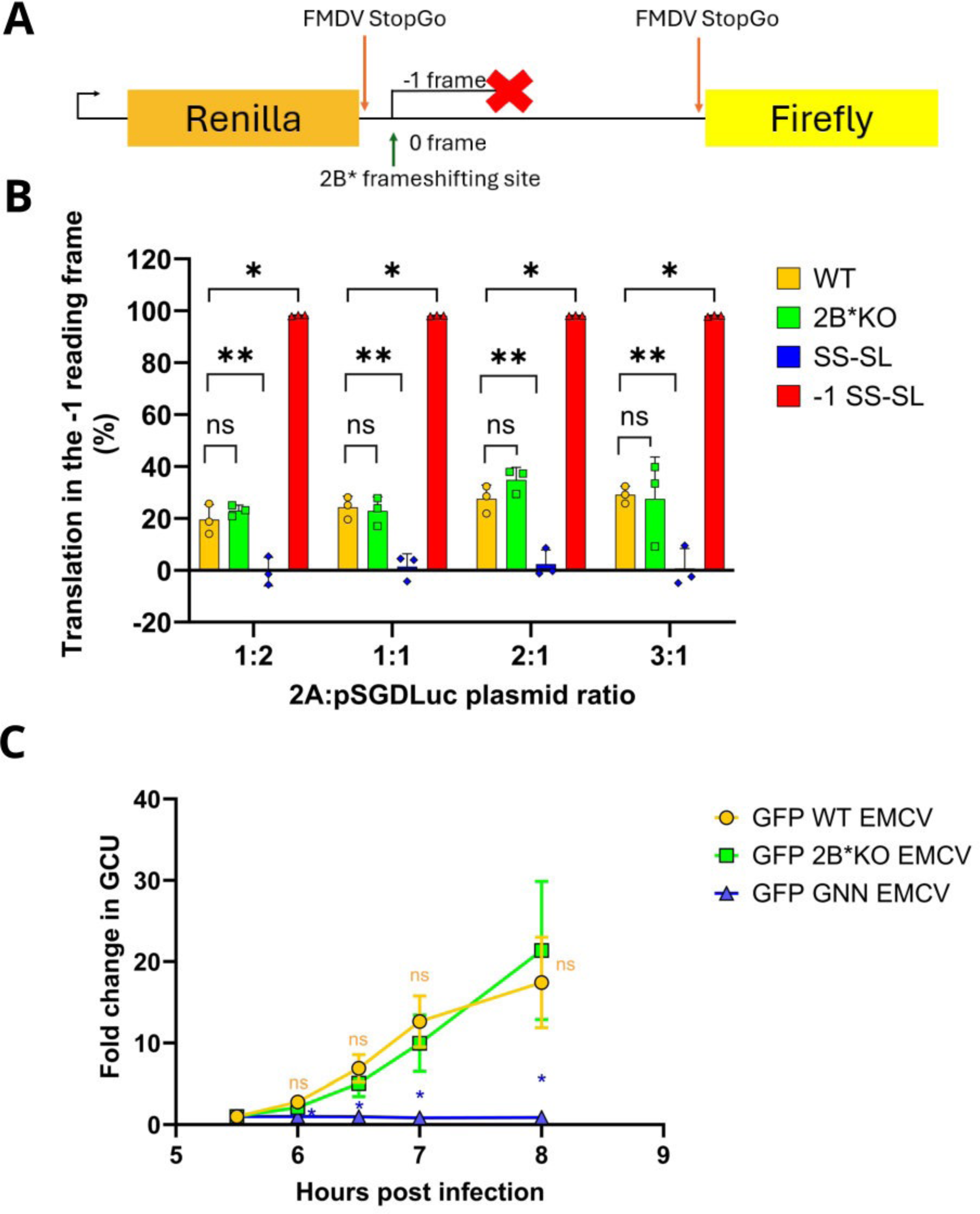
2B*KO EMCV mutations do not impact viral frameshifting or replication. **(A)** Schematic representation of the dual luciferase constructs. Renilla luciferase is produced by every translating ribosome. However, in the WT sequence, firefly luciferase is only produced when frameshifting does not occur. This enables quantitative measurement of the percentage of ribosomes in each reading frame for each mutant construct. **(B)** BHK-21 cells were transiently co-transfected with a WT EMCV 2A expression construct (pCAGG-2A) along with a dual luciferase plasmid containing 105 nt of the EMCV genome (WT pSGDLuc) or mutants thereof (2B*KO pSGDLuc, SS-SL pSGDLuc or −1 SS-SL pSGDLuc) at various ratios. At 24 h post-transfection, cells were frozen in 1x PLB and both renilla and firefly luciferase activities were measured. Samples were normalised to the luciferase values for the same pSGDLuc construct co-transfected with pCAGG-2Amut. The percentages of ribosomes in the 0 or −1 reading frames for each ratio of pCAGG-2A:pSGDLuc were calculated. Data shown are the mean +/− SD of three biological repeats, each using triplicate wells. Statistical analysis (Student’s *t*-test): ns = not significant, ∗ *p* ≤ 0.05, ∗∗ *p* ≤ 0.01. **(C)** A confluent monolayer of BHK-21 cells was infected with GFP WT EMCV or the indicated mutants at MOI 0.01. GCU was analysed using the Incuyte live-cell imaging suite (Sartorius) and normalised to respective GCU at 5.5 h p.i. Data shown are the mean +/− SD of three biological repeats, each using triplicate wells. Statistical analysis (Student’s t-test of the indicated virus compared to GFP WT EMCV): ns = not significant, * *p* ≤ 0.05.

For each sequence tested, the shift site, stem loop and any downstream mutations were inserted into the previously described pSGDLuc plasmid (12). The resulting plasmids contained 11 nt upstream of the slippery sequence, the 7 nt slippery sequence itself and an additional downstream 87 nt, between renilla and firefly luciferase ORFs which were in the same reading frame (0 frame) (Figure 1A). Flanking the inserted nucleotides were two FMDV StopGo sequences, enabling co-translational separation, to ensure that the peptide sequences encoded by the insert are not tagged onto the end of the renilla and/or firefly luciferases where they might differentially affect enzymatic activities. Ribosomes which do not frameshift translate both luciferase proteins whereas riboosomes which frameshift translate only the renilla luciferase (Figure 1A).

As binding of the EMCV 2A protein to the stem-loop is necessary for PRF, a construct encoding FLAG-tagged EMCV 2A (pCAGG-2A) was co-transfected with each pSGDLuc construct. To allow normalisation of the relative luciferase readings, each pSGDLuc construct was also co-transfected with a plasmid encoding a FLAG-tagged 2A mutant (pCAGG-2Amut) that is unable to bind the stem-loop and unable to stimulate PRF (3). The PRF efficiency could then be calculated as 1 − (FLuc_test_/RLuc_test_)/(FLuc_2Amut_/RLuc_2Amut_). As the ratio of pCAGG-2A to pSGDLuc which would mimic the ratio of 2A protein to viral RNA occurring in infection was unknown, a range of co-transfection ratios were performed (Figure 1B).

A previously described mutant sequence, non-permissive to PRF due to mutations in both the slippery sequence and stem-loop (SS-SL) (3), was also inserted in the pSGDLuc vector as an additional control. A further single-nucleotide insertion was added to this construct to create −1 SS-SL. Here, the SS-SL mutations prevented PRF and the insertion caused all ribosomes to enter the 2B*-derived reading frame, thus mimicking 100% PRF. As expected, pSGDLuc-2B*KO showed no significant difference in PRF efficiency compared to pSGDLuc-WT, indicating that the 2B*KO mutations do not affect PRF and therefore any phenotypic differences found between 2B*KO EMCV and WT EMCV in later experiments would not be due to impaired PRF efficiency.

In principle, any change to the viral RNA genome may also influence viral RNA replication. 2B* mutant viruses have previously been observed to reach equivalent overall titres to WT (2). Furthermore, 2B* is only expressed at late time points, after substantial amounts of RNA replication have already occurred. Therefore any difference in viral replication rates between 2B*KO EMCV and WT EMCV would likely be due to changes in RNA sequence or RNA structure, rather than a direct function of the 2B* protein. To further validate the use of 2B*KO EMCV as a tool for later studies, we investigated the effect of the 2B*KO mutations on viral replication using a GFP-tagged virus (GFP WT EMCV) (13), which contains the GFP ORF immediately followed by the EMCV 3C protease cleavage site, upstream of the leader and capsid proteins. The essential GDD motif in the viral RNA polymerase, 3D, was mutated to GNN to create a replication incompetent control. The GNN mutation and the 2B*KO mutation were independently introduced into the parental genome, creating GFP GNN EMCV and GFP 2B*KO EMCV, respectively. Following infection, the increasing levels of GFP in an infected cell would be directly proportional to the amount of viral protein produced and hence a reflection of viral replication.

BHK-21 cells were infected with these viruses and viral replication over time was measured by live-cell imaging (Figure 1C). Green calibrated units (GCU) at each timepoint were normalised to the GCU in that sample at 5.5 h p.i., which was when GFP was first detectable in most samples. As cell lysis occurred at approximately 8 h p.i. (one viral replication cycle), GFP was not measured past this point. The fold change in GCU was not significantly different between cells infected with GFP WT EMCV or GFP 2B*KO EMCV at any of the timepoints measured (Figure 1C) while GFP GNN EMCV consistently gave only background levels of GCU. Thus the 2B*KO mutations do not appear to affect viral replication. Therefore, any differences found between 2B*KO EMCV and WT EMCV are unlikely to be caused by defects in either PRF or viral replication and are likely to be solely attributable to the loss of 2B* protein.

### Characterisation of a virus expressing HA-tagged 2B*

Next we modified the EMCV genome to encode 2B* with an N-terminal HA tag (HA2B* EMCV). This would allow 2B* to be immunoprecipitated from infected samples, and the subsequent identification of its binding partners. The HA tag and a short Gly-Ser flexible linker was encoded immediately following the final proline of the StopGo sequence, responsible for separating 2A from 2B/2B* (Figure 2A). A similar approach has been previously described to tag 2B* with a V5 epitope (11). As 2B and 2B* share the first 12 residues prior to the change in reading frame, both 2B and 2B* will be translated with the N-terminal HA tag. However, the viral 3C protease is thought to cleave the N terminus of 2B, thereby removing the tag (1, 11). The KO mutations were also introduced into HA2B* EMCV to create an equivalent KO mutant virus.

**Figure 2:**
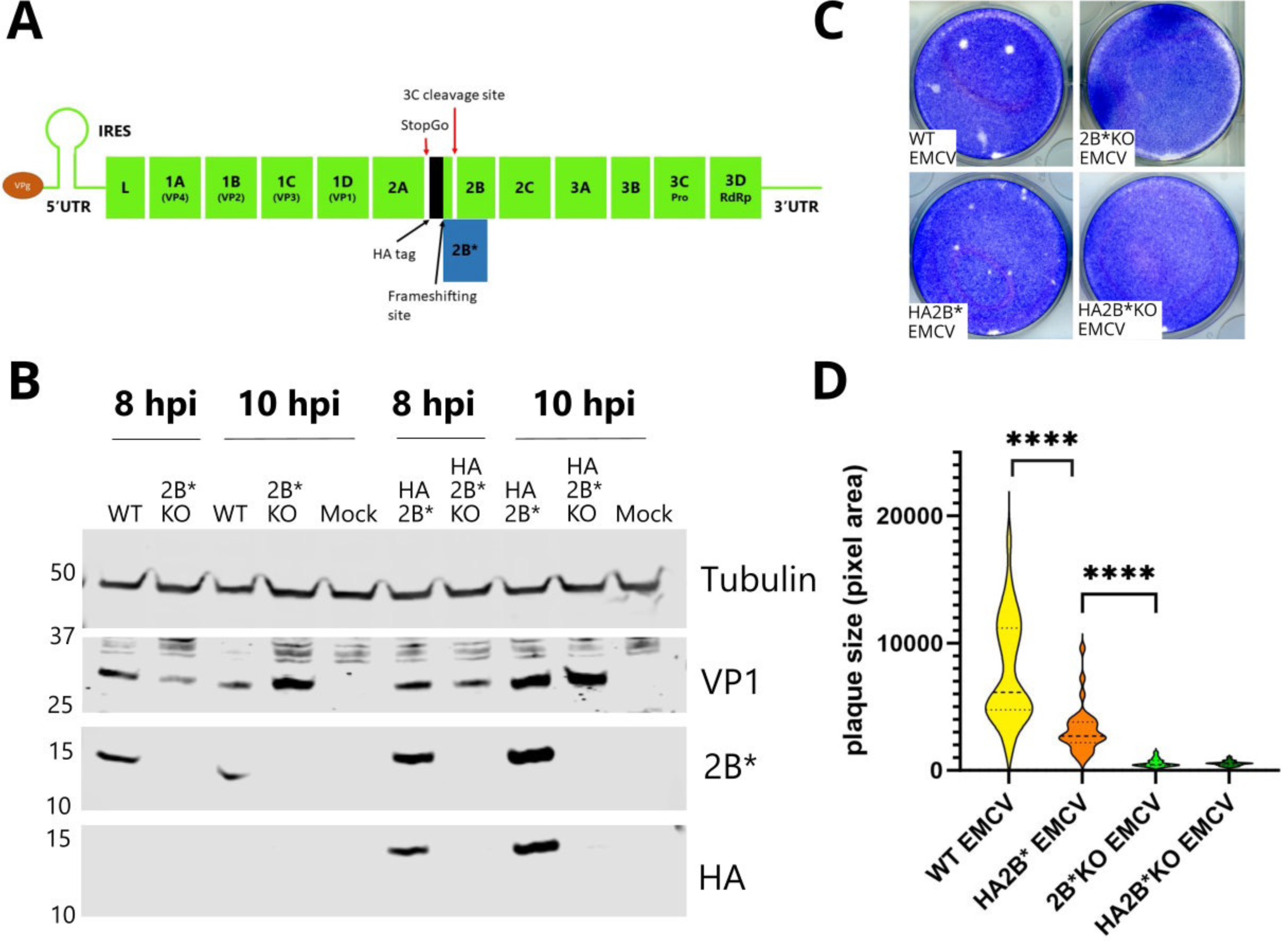
Characterisation of HA2B* EMCV. **(A)** Schematic representation of the EMCV viral genome with sequence encoding the HA tag inserted to tag 2B*. **(B)** BHK-21 cells were infected with WT EMCV, 2B*KO EMCV, HA2B* EMCV or HA2B*KO EMCV at MOI 5.0. Cells were harvested at the indicated time points, and frozen in complete RIPA buffer. Lysates were subjected to SDS-PAGE and immunoblotting. The membrane was cut at the 25 kDa marker on the ladder to allow the same membrane to be probed by all four antibodies. The molecular mass scale (kDa) is indicated at left and antibodies used are labelled at right. **(C)** Images of plaques formed by the viruses indicated, grown in BSR cells. Images are representative of three independent biological repeats. (**D)** Size distribution of plaques formed by WT EMCV, HA2B* EMCV, and their respective 2B*KO mutants. Distributions shown are based on area measurements of 30 plaques, sampled from three biological repeats. Horizontal lines represent the median (dashed) and upper and lower quartiles (dotted). Statistical analysis (ratio-paired *t*-test): ∗∗∗∗ *p* ≤ 0.0001.

At both 8 and 10 h p.i., HA2B* EMCV produced detectable levels of HA-tagged 2B* in BHK-21 cells (Figure 2B). Noticeably, HA-tagged 2B was not detected in any of the lysates (predicted molecular mass 13 kDa), indicating that the HA epitope was indeed cleaved from the majority of 2B as expected.

To ensure that the addition of the HA tag did not disable the function of 2B* and that HA2B* KO EMCV recreated the small plaque phenotype associated with the loss of 2B* (2, 3), the plaques produced by each virus at 48 h p.i. in BSR cells were measured (Figure 2C, 2D). Although HA2B* EMCV created smaller plaques than untagged WT EMCV (median areas 2694 pixels and 6140 pixels, respectively) (Figure 2D), they were still significantly larger than those of 2B*KO EMCV (median area 451 pixels). The decreased size of HA2B* EMCV plaques indicated that the addition of the tag did somewhat affect 2B* function. However this change was modest and, as the plaques were still substantially larger than those created by 2B*KO EMCV, it is likely that HA-tagged 2B* still interacts with the relevant factors required for the large plaque phenotype. Therefore, HA2B* EMCV was used as a tool to characterise the interaction partners of 2B*.

### Mass spectrometry identifies 14-3-3 protein family members as putative interaction partners of 2B*

To identify the host and viral protein interactions of 2B*, HA2B* was immunoprecipitated from HA2B* EMCV infected BHK-21 cells at 7 h p.i. in triplicate. Immunoprecipitated samples were then subjected to trypsin digestion and labelling by tandem-mass-tagging (TMT) prior to being analysed by liquid chromatography with tandem mass spectrometry (LC-MS/MS) (Figure 3A). While the predominant tagged viral protein was 2B*, the StopGo mechanism is approximately 94% efficient in EMCV and therefore small amounts of HA-tagged 2A-fusion products (2AHA2B* and 2AHA2B^N^, where 2B^N^ represents the N-terminal fragment of 2B upstream of the 3C protease cleavage site (11) were produced (Figure 3B). To ensure any interacting proteins bound to these fusion products could be identified and removed from analysis, HA2B*KO EMCV infected controls were included in addition to mock infected controls.

**Figure 3:**
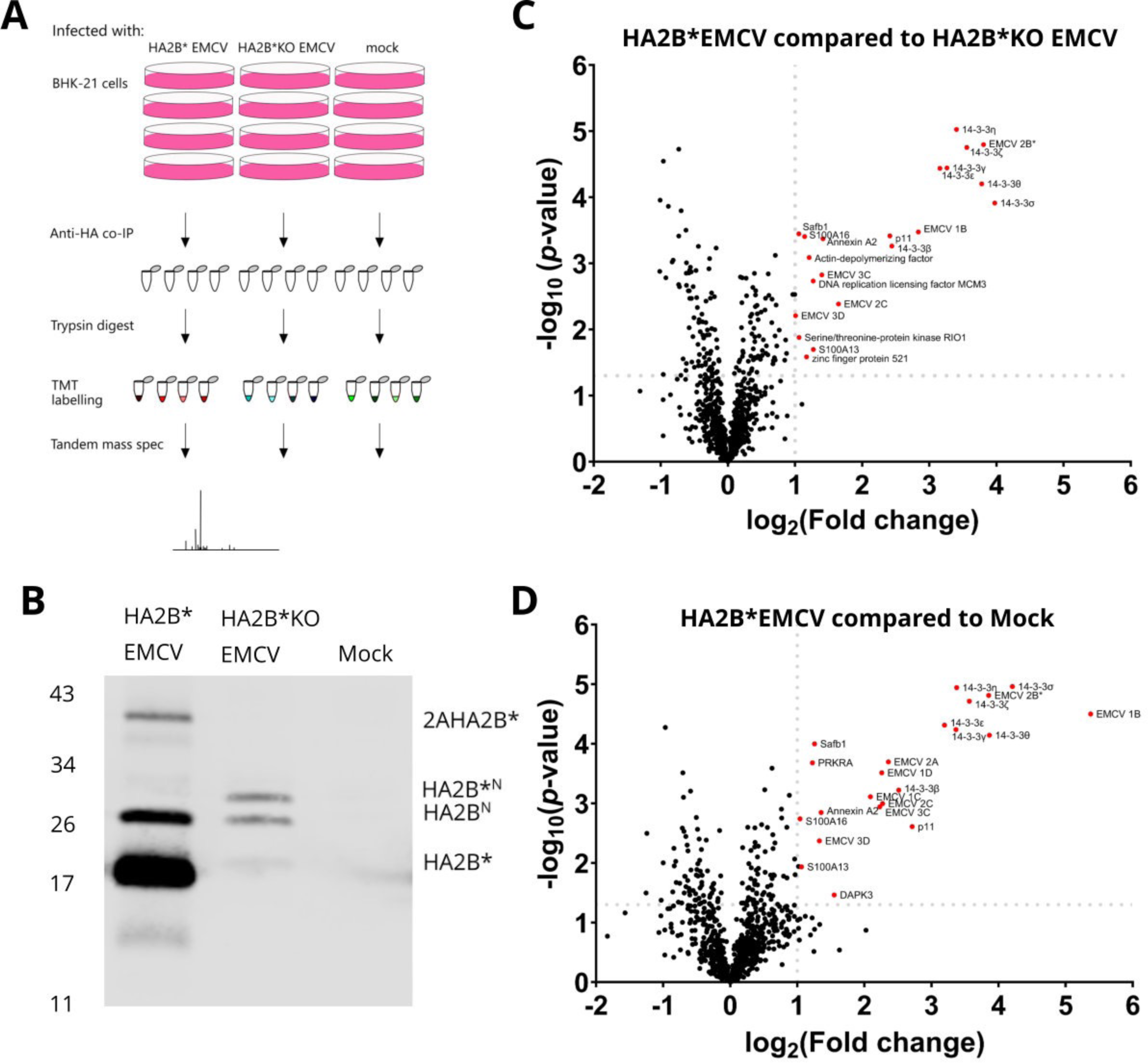
Putative interaction partners of 2B* include the entire family of 14-3-3 proteins. **(A**) Schematic diagram of the TMT experiment. BHK-21 cells were infected with either HA2B* EMCV or HA2B*KO EMCV, or mock infected. At 7 h p.i., HA2B* along with any interaction partners were purified from the lysates by anti-HA co-immunoprecipitation; 87.5% of each sample was then used for trypsin digestion, TMT labelling and analysis by mass spectrometry. **(B**) The remaining 12.5% of each co-immunoprecipitated sample was probed for HA-tagged proteins by western blot using an anti-HA antibody. The molecular mass scale (kDa) is indicated at left and HA-tagged proteins are labelled at right. (**C**) and (**D**) Two-sample *t*-tests were used to compare protein enrichment in the HA2B* EMCV infected samples relative to HA2B*KO EMCV infected samples (panel C) and mock infected samples (panel D). The −log_10_(*p*-value) greater than 1.3 (*p*-value >0.05) and fold change of greater than 2 (log_2_(Fold change > 1) for each recognised protein are shown. Candidate interaction partners were defined as those with both a Difference greater than 1 and a −log_2_ (*p*-value) greater than 1.3. Only proteins which were above these thresholds in the specified comparison are labelled on the graph (red).

Seventeen proteins were enriched in HA2B* EMCV infected samples relative to both HA2B* KO EMCV infected and mock infected samples, including HA2B* itself (Table 1, Figure 3C, 3D). Candidate interaction partners were defined as those having both a Student’s *t*-test difference greater than 1.0 and a −log_2_(*p*-value) greater than 1.3. Interestingly, this included the entire family of 14-3-3 proteins. The 14-3-3 family are scaffold proteins involved in a wide range of cellular functions, including apoptosis (14, 15), cell cycle progression (16, 17), nuclear trafficking (18), innate immune signalling (19, 20), autophagy (21) and proteasome function (22). Due to the importance of these proteins in many pathways which EMCV utilises, we chose to focus on these proteins for subsequent analyses.

To confirm their interaction with 2B*, a construct expressing HA2B* was co-transfected into BHK-21 cells along with each construct encoding an N-terminally FLAG-tagged 14-3-3 protein, cloned from BHK-21 cDNA. As 14-3-3ε isoforms X1 and X2 could not be distinguished by the peptides detected by mass spectrometry, both were included. The 14-3-3 proteins were then immunoprecipitated from the samples via the FLAG-tag. Immunoprecipitated samples were subjected to SDS-PAGE and immunoblotting to determine whether HA2B* was bound to these putative interaction partners (Figure 4A). As tubulin β-chain was not identified by the co-immunoprecipitation mass spectrometry analysis as a potential interaction partner of HA2B*, a construct encoding N-terminally FLAG-tagged tubulin β-chain (FLAG-TUBB) was included as a negative control.

**Figure 4:**
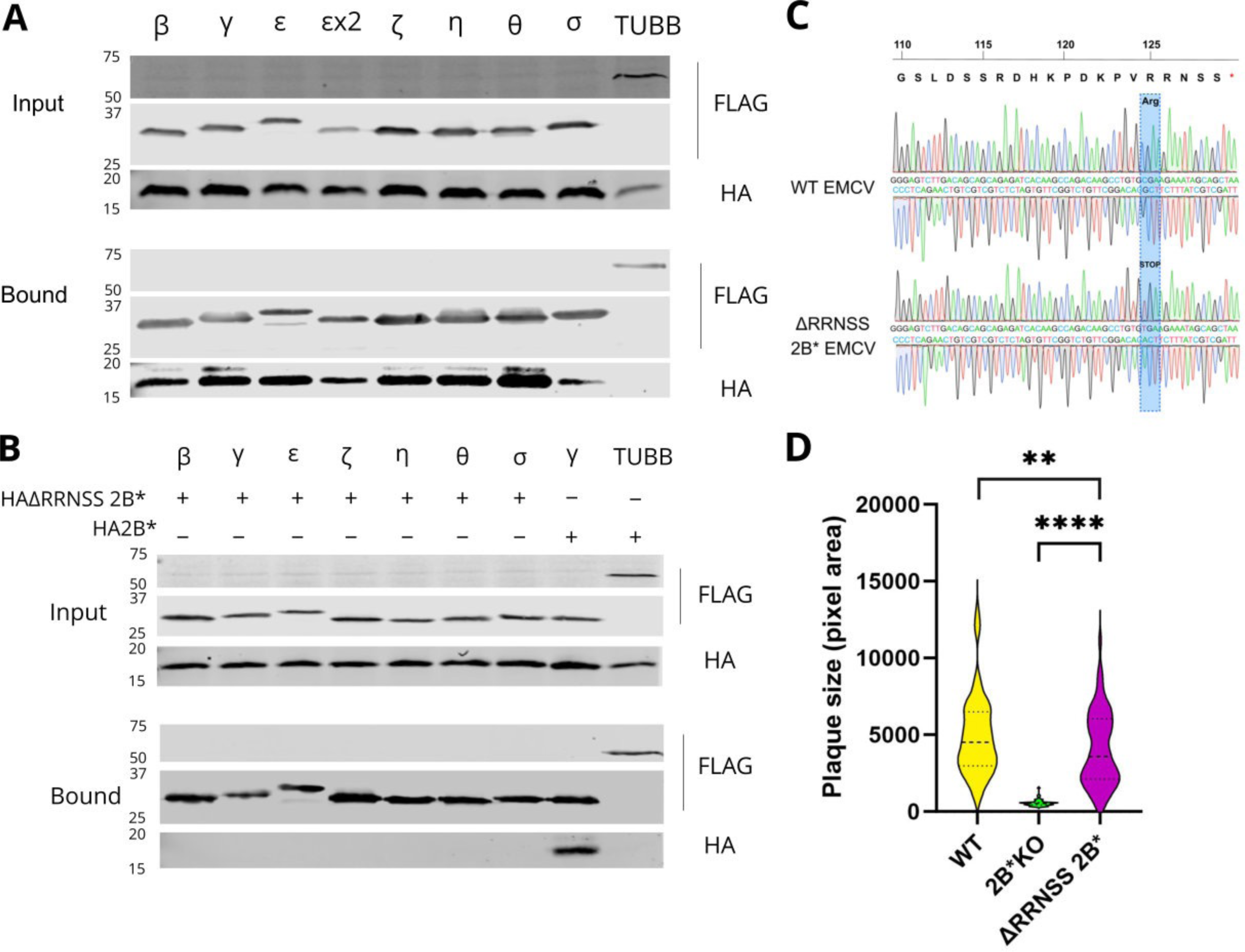
2B* binds to the entire family of 14-3-3 proteins via a C-terminal RRNSS sequence. **(A)** BHK-21 cells were co-transfected with equal amounts of pCAGG-HA2B* and the specified N-terminally FLAG-tagged 14-3-3 encoding plasmid (pCAGG-FLAG 14-3-3x), 24 h prior to immunoprecipitation via the FLAG epitope. Samples were then subjected to SDS-PAGE and immunoblotting using the indicated antibodies. Data shown are representative of two independent biological repeats. **(B)** BHK-21 cells were co-transfected with both a plasmid encoding either an N-terminally FLAG-tagged 14-3-3 protein or tubulin β chain and another encoding either HAΔRRNSS2B* or HA2B*, as indicated. Cell lysates were immunoprecipitated with anti-FLAG antibody and subjected to SDS-PAGE and immunoblotting. Data shown are representative of two independent biological repeats. **(C)** Confluent monolayers of BHK-21 cells were infected with WT EMCV or ΔRRNSS 2B* EMCV at MOI 0.01. At 24 h p.i., RNA was extracted from each sample and subjected to RT-PCR to amplify the region of interest. The cDNA was sequenced (Sanger method) with both forward and reverse primers. Both chromatograms have only one clear peak for each nucleotide, indicating each infected sample contained only one EMCV sequence detectable by Sanger sequencing. Chromatograms shown are representative of three independent biological repeats. **(D)** BSR cells were infected with the indicated viruses 1 h prior to being overlaid with semi-solid medium (1% LMA) for 48 h. Distributions shown are based on area measurements of 60 randomly chosen plaques sampled over three biological repeats. Horizontal lines represent the median (dashed) and upper and lower quartiles (dotted). Statistical analysis (ratio-paired *t*-test): ∗∗ *p* ≤ 0.01, ∗∗∗∗ *p* ≤ 0.0001.

The interaction between 2B* and all of the 14-3-3 proteins was verified as the FLAG-tagged proteins were all found to co-precipitate HA2B* (Figure 4A). These proteins are therefore able to bind to 2B* both during infection and in overexpression and thus other viral proteins are not required to mediate this interaction. Confocal microscopy confirmed that 2B* and all 14-3-3 family members were dispersed throughout the cytosol (Figures S1, S2, S3), and redistribution was not observed during co-expression.

### 14-3-3 binding is mediated by a C-terminal 2B* motif and is not responsible for the 2B*KO small plaque phenotype

To identify the motif(s) within 2B* responsible 14-3-3 binding, we utilised the online resource ELM (eukaryotic linear motifs) (23), which searches for short linear motifs corresponding to known protein interaction sites. The ELM tool predicted the C-terminal RRNSS sequence of 2B* to be a binding site for the 14-3-3 protein family. Indeed, this was the best-scoring potential interaction site in 2B*, with a reported *p*-value of 6.4 × 10^−5^. The C-terminal RRNSS sequence is highly conserved across EMCV strains with occasional variations in the last position (to N, I or L). The known 14-3-3 binding motif is highly conserved and, when at the C terminus of the binding partner in question, is invariably p[S/T]-X_1–2_-COOH with upstream arginine residues preferred (RRXp[S/T]-X_1–2_-COOH) (where p is one or more phosphate group) (24). The 14-3-3 proteins form homo-and heterodimers which bind target proteins when the penultimate serine within the recognition motif is phosphorylated (24, 25), leading to translocation or sequestration of the binding partner in question.

To investigate whether this sequence is indeed required for 2B*:14-3-3 binding, an RRNSS truncated HA-tagged 2B* construct (HAΔRRNSS 2B*) was engineered using a R125Stop mutation, thus removing the five C-terminal residues of 2B*. This construct was co-transfected into BHK-21 cells alongside each construct encoding a FLAG-tagged 14-3-3 protein, and the FLAG co-immunoprecipitation assay was repeated (Figure 4B). Every FLAG-tagged member of the 14-3-3 family was entirely unable to bind to the co-transfected ΔRRNSS2B* mutant whereas the WT HA2B* was still successfully bound by a representative 14-3-3 protein, 14-3-3γ. This clearly demonstrates that the RRNSS sequence of 2B* is essential for the 2B*:14-3-3 interaction and deletion of this sequence completely prevents this interaction.

We next investigated whether the lack of the 14-3-3:2B* interaction was the cause of the small plaque phenotype characteristic of 2B* KO viruses (2, 3). The R125Stop mutation, which is synonymous in the 2B reading frame, was introduced into the viral genome, creating ΔRRNSS 2B* EMCV. To confirm that this mutation did not revert during infection, the virus produced from infected BHK-21 cells was sequenced by Sanger sequencing (Figure 4C). No detectable reversion was observed by 24 h p.i.

Interestingly, the small plaque phenotype associated with 2B*KO EMCV (median plaque area 524 pixels) was not reproduced by ΔRRNSS 2B* EMCV (Figure 4D). Although the difference between WT EMCV and ΔRRNSS 2B* EMCV plaque sizes was still statistically significant, it was very modest (median areas 4516 pixels and 3577 pixels, respectively). Therefore, the 2B*:14-3-3 interaction is unlikely to be responsible for the increased plaque size seen in WT EMCV infection.

Presumably, the 2B*:14-3-3 interaction performs an additional function that is beneficial to the virus. We hypothesised that the sequestration of 14-3-3 proteins by 2B* may contribute to the evasion of innate immune signalling as other viral binding partners of 14-3-3 proteins – such as the 3C protease of enterovirus – are known to have similar effects (7–10). During viral infection, RIG-I is relocalised from the cytosol to the mitochondria by 14-3-3ε which forms a complex with both RIG-I and TRIM25 (20), the ubiquitin ligase essential for the antiviral function of RIG-I (26), enabling the activation of MAVS on the mitochondrial membrane. In addition, 14-3-3η is responsible for the redistribution of MDA5, enabling MDA5-induced antiviral IFN-β signalling via MAVS (19). As each 14-3-3 monomer has only one binding pocket, it is unlikely that 14-3-3 proteins bound to 2B* would be able to perform these functions. Therefore, we next investigated the ability of 2B* to antagonise innate immune signalling via the 14-3-3 interaction.

### The 2B*:14-3-3 interaction interferes with innate immune signalling outside of the context of infection

MEF cells were transiently transfected with pCAGG-HA2B*, pCAGG-HAΔRRNSS2B* or empty pCAGG 24 h prior to activation of the IRF3 and NFκB signalling pathways by transfection with poly(I:C), a dsRNA mimic. After 6 h, *IFNB1*, *ISG15*, *IL6*, *RSAD2* and *IFIT1* transcript levels were quantified by qRT-PCR, normalised to GAPDH mRNA using the ΔΔCT method, and compared to those for the empty pCAGG transfected samples. While overexpression of HA2B* prior to poly(I:C) stimulation significantly reduced the transcription of *IFNB1*, *IL6* and the ISGs *ISG15* and *RSAD2* involved in innate immune signalling (Figure 5A), this antagonistic ability was abrogated by the loss of the RRNSS motif responsible for 14-3-3 binding.

**Figure 5:**
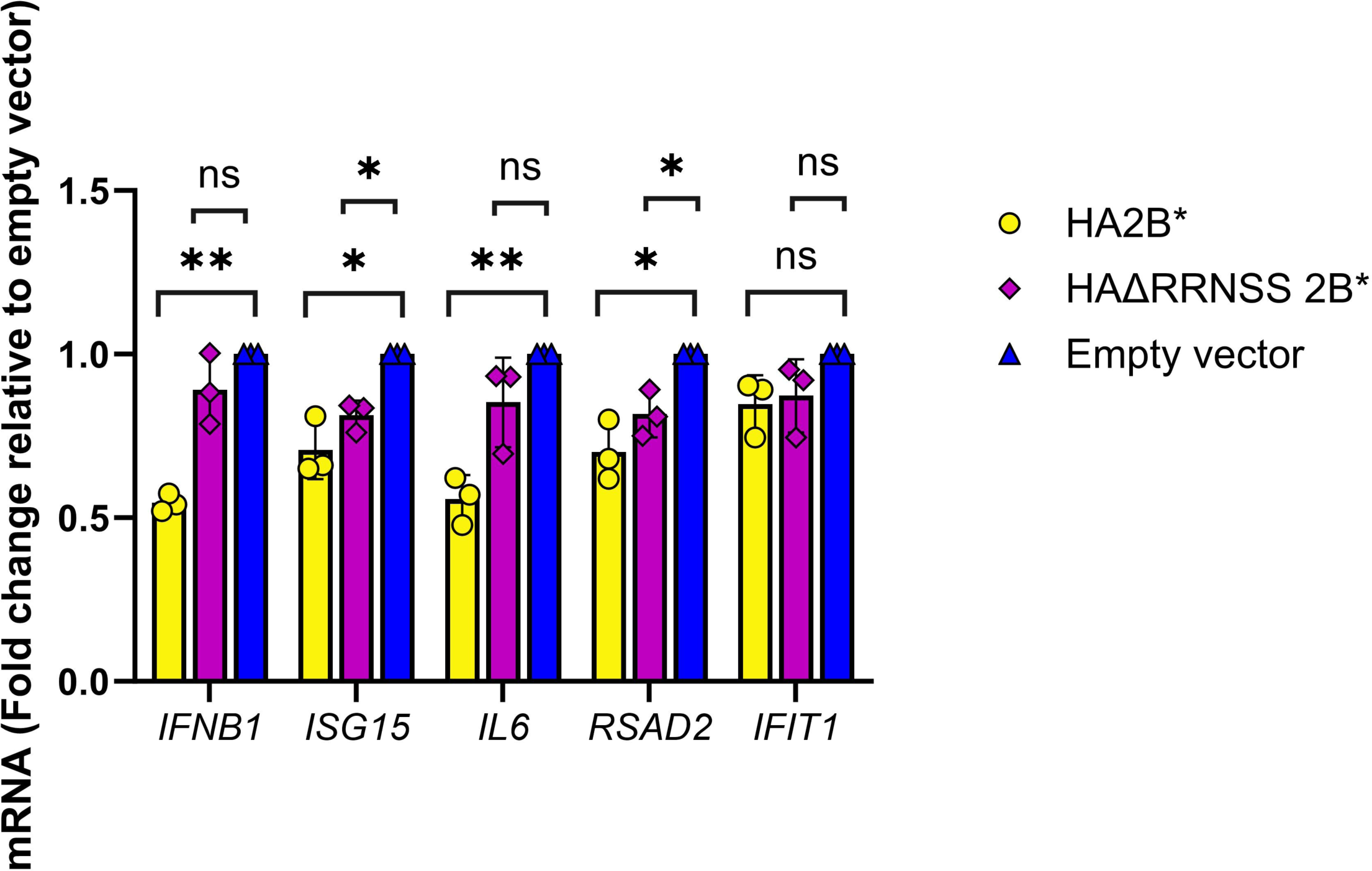
Overexpressed 2B* interferes with innate immune signalling via its interaction with 14-3-3 proteins. **(A)** MEF cells were transiently transfected with constructs encoding HA2B* or HAΔRRNSS 2B*, or the empty vector, as indicated 24 h prior to transfection with poly(I:C) (final concentration 10 ng/mL) for 6 h. The relative expression level of each gene was determined by qRT-PCR. Expression levels were normalised internally to GAPDH and to poly(I:C) stimulated samples transfected with the empty vector (pCAGG). Data shown are the mean +/− SD of three biological repeats. Statistical analysis (unpaired Welch *t*-test): ns = not significant, ∗ *p* ≤ 0.05, ∗∗ *p* ≤ 0.01.

## DISCUSSION

In this study, we successfully engineered an EMCV genome which encodes an HA-tagged 2B* protein, enabling the first characterisation of the binding partners of 2B*. Whilst the addition of the tag had a modest effect upon plaque size, it did not completely abrogate 2B* function (Figure 2C, 2D). We also confirmed that the 2B*KO virus had WT levels of PRF and RNA replication, providing future studies with a tool for studying 2B* phenotypes without the problem of confounding effects (Figure 1B, 1C). We identified 17 putative binding partners of 2B*, seven of which are members of the 14-3-3 family, and identified the C-terminal 5 residues of 2B* as being essential for the 2B*:14-3-3 interaction. Finally, we determined the functional consequence of this interaction to be disruption of innate immune signalling, presumably through sequestration of 14-3-3 proteins preventing their contribution to MDA5 and RIG-I activation (19, 20) and subsequent inflammatory cytokine induction.

Upon recognition of viral RNA, MDA5 and RIG-I translocate from the cytosol to the mitochondria where they activate MAVS (27), leading to NFκB and IRF3 signalling (28) and the production of antiviral cytokines including IL-6 and IFN-β, respectively. The mechanisms by which picornaviruses antagonise this type I IFN signalling pathway vary widely, and most species utilise multiple synergistic mechanisms exerted by different viral proteins. It is therefore unsurprising that 2B* has evolved to reduce the immune response to EMCV infection, despite EMCV encoding multiple other proteins with innate immune inhibitory functions (namely, 2C, 3C and L). Although overexpression of 2B* resulted in significant reduction of *IFNB1* and *IL6* transcription (Figure 5A), we were unable to detect these transcripts by qRT-PCR in innate-competent RAW264.7 cells infected with either WT EMCV or ΔRRNSS 2B* EMCV (data not shown). A possible explanation is that – at least in these cells – other EMCV proteins compensate for the loss of this function of 2B* and still prevent large amounts of *IL6* and *IFNB1* mRNA from being produced.

To reduce recognition of dsRNA, EMCV viral protein 2C directly interacts with MDA5, limiting MDA5-mediated innate immune signalling during EMCV infection (29). MDA5 signalling has been suggested to be further impaired during EMCV infection by EMCV capsid protein VP2 (23), although the relevance of this role has yet to be confirmed in virus infection. The EMCV protease 3C directly cleaves both RIG-I and TANK, further limiting dsRNA recognition and promoting NFκB signalling over IRF3 dimerisation. Converging upon this effect, EMCV protein L impedes transcriptional upregulation of antiviral response genes both by preventing IRF3 dimerisation and by inhibiting nucleocytoplasmic trafficking of the relevant transcription factors (30). However, this is not true for all cardioviruses: the L protein from some TMEV strains reduces IFN-β production but only after the dimerised IRF3 has entered the nucleus (31). In addition, the TMEV-specific L* protein further reduces the type I IFN response by binding to and preventing the dimerisation of RNase L, preventing IRF3 activation (32, 33). The 14-3-3 sequestration mechanism exhibited by the 2B* protein appears distinct from these previously described mechanisms in cardioviruses and therefore provides an additional layer of protection for the virus from the host response.

As the presence of 2B* appeared to reduce the levels of all antiviral transcripts tested (Figure 5A), we cannot eliminate the possibility that the 2B*:14-3-3 interaction causes a global reduction in transcription initiation instead of specifically influencing the antiviral response. However, 2B* does not have a nuclear localisation (Figure S1, S2, S3) and, although the 14-3-3 proteins are involved in nuclear transport of some transcription factors (18, 34), these do not include IRF3 or NFκB which are responsible for the upregulation of the target genes tested (*IL6, IFNB1, ISG15, IFIT1* and *RSAD2*). In addition, EMCV-induced reduction of host transcription and translation (so-called host shutoff) occurs from 4 h p.i., before 2B* is efficiently translated (11, 35). Therefore, it is most likely that the effect is mediated prior to the nuclear import of IRF3 and NFκB, as the 2B*:14-3-3 interaction would impede the earlier translocation of both RIG-I and TRIM25 by 14-3-3ε (20) and MDA5 by 14-3-3η (19) (Figure 6), in a manner similar to the mechanisms exerted by the 14-3-3-binding proteins of influenza A, enteroviruses, Zika virus and Epstein-Barr virus (7–10).

**Figure 6:**
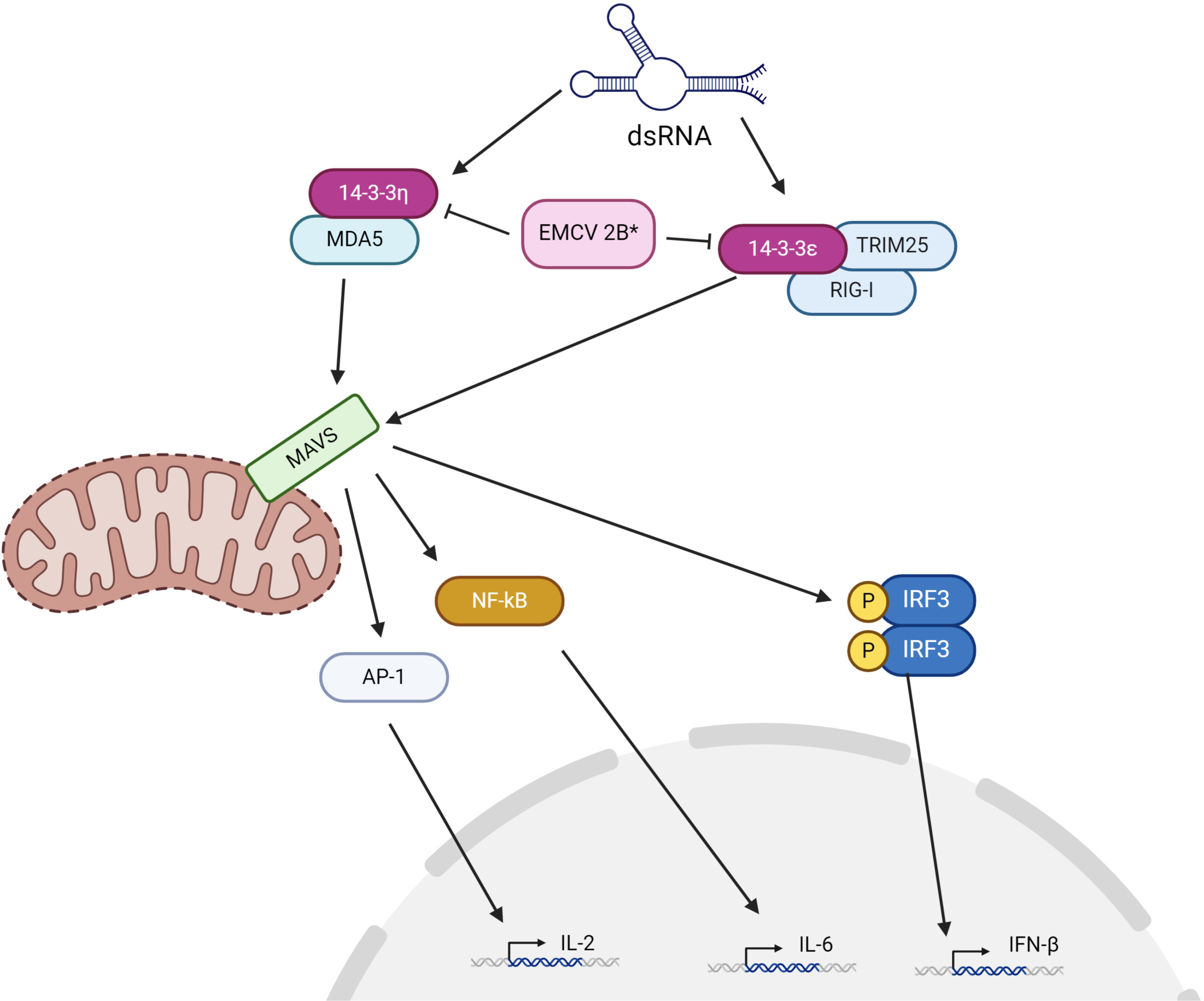
Model of 2B* function. 2B* reduces NFκB and IRF3 signalling likely by inhibiting the translocation of innate immune signalling molecules MDA5, RIG-I and TRIM25 by preventing their interactions with 14-3-3 proteins, thus reducing MAVS activation.

The 14-3-3 proteins are a family of scaffold proteins which form both homo-and heterodimers, performing a plethora of roles many of which are unrelated to antiviral immunity; hence many as-yet-undiscovered pathways may also be affected by 14-3-3 interactions with 2B*. The 14-3-3 dimeric complex can bring two 14-3-3-bound proteins into proximity, to increase the chances of interaction. Conversely, they may prevent interactions by sequestering proteins away from alternative interaction partners. Although the binding motif is relatively conserved between all 14-3-3 proteins, each also have distinct interaction partners to enable execution of specific roles. Although 2B* was found to interact with the entire family of 14-3-3 proteins in our study, the relative binding affinity of 2B* to each family member has not yet been investigated. It is also possible that some of these interactions are indirect and are mediated by the formation of 14-3-3 heterodimers. Further investigation may uncover specificity in the 2B* interaction, perhaps ensuring 2B* preferentially affects particular pathways *in vivo*. Binding is controlled by the phosphorylation status of the target protein, as the recognition motif is only bound once phosphorylated(14); the relevance of phosphorylation for 2B* function has also not yet been investigated. The 14-3-3 proteins also preferentially bind to disordered regions (36) (which the C-terminus of 2B* is predicted to be by analysis with Alphafold, I-TASSER and NetSurfp).

It seems counterintuitive that 2B*, a protein which is only translated at late timepoints shortly before the onset of cell lysis, would have a role in preventing the antiviral response. However other viruses are known to add further layers of protection against innate immune signalling at later stages of infection (37–39). This is likely due to the additional need for protection following the increased pathogen associated molecular patterns (PAMP) exposure caused by cell death and permeabilisation of the membranes: the change in cytosolic composition also induces innate immune responses (40), hence there is increased risk of forewarning surrounding cells of virus exposure during the lysis process. In addition, at early stages of infection EMCV benefits from NFκB activation (41) which would no longer be necessary when lysis is underway. It therefore seems reasonable that 2B* could provide an additional layer of protection against antiviral responses at this late stage. However, since TMEV also contains the PRF signal, but not the lengthy 2B* ORF, it is likely that highly efficient temporally controlled PRF evolved first as a ribosome sink to downregulate synthesis of the replication proteins at late timepoints. Expansion of the 2B* ORF in the EMCV lineage allowed 2B* to secondarily evolve into a functional protein in its own right. Thus, late expression of 2B* may alternatively be an unavoidable consequence of the original function of cardiovirus PRF and is not necessarily optimal for its role in antagonising innate immune signalling.

A focus of future work will be to assess the function of the other binding partners identified in our proteomics screen; it is possible that at least one of these proteins contributes to the small plaque phenotype already described. The minor reduction in plaque size produced by ΔRRNSS 2B* EMCV (Figure 4D) may potentially be explained by a slight increase in MAVS-induced apoptosis or MAVS-induced inhibition of viral spread, and clearly this interaction does not fully explain the plaque size phenotype previously reported for 2B*KO viruses (2, 3).

In summary, we propose a model whereby 2B* inhibits translocation of the innate immune signalling molecules MDA5, RIG-I and TRIM25 via its interaction with 14-3-3 proteins, thereby reducing NFκB and IRF3 signalling. Our findings add 2B* to the growing list of viral proteins, from a wide range of families, which interact with the 14-3-3 proteins to reduce antiviral cytokine signalling (7–10) and identify 2B* as another “transframe” protein with a role in antagonising the innate immune response (37, 42, 43).

## MATERIALS AND METHODS

### Mammalian cell culture

BHK-21 (ATCC) (baby hamster kidney fibroblast), BSR (single cell clone of BHK-21 cells) and MEF (mouse embryonic fibroblast) cells were maintained in Dulbecco’s Modified Eagle Medium with high glucose (Sigma), supplemented with 10% (v/v) heat-inactivated foetal calf serum (FCS), 25 mM HEPES, 2 mM L-glutamine and non-essential amino acids (Sigma) (“complete DMEM") in a humidified 5% CO_2_ atmosphere at 37°C. All cell lines were confirmed to be mycoplasma-free at regular intervals (MycoAlert^TM^ PLUS Assay, Lonza).

### RT-PCR

Uninfected BHK-21 cells, or cells infected at an MOI of 0.1, were trypsinised either 24 h after plating or 24 h p.i. respectively. The trypsin was neutralised in 10% FCS DMEM before the cells were pelleted at 1000 × *g* and the media was removed before the RNA was extracted using a commercial kit as recommended (RNeasy, Qiagen). Reverse transcription of each sample was carried out using 5 µg of RNA, random hexamers and Superscript III reverse transcriptase (ThermoFisher Scientific). PCR using relevant primers immediately followed reverse transcription.

### Expression Plasmids

The pCAGG-HA2B* expression construct was originally synthesised from a gene block (Integrated DNA Technologies), designed with the 2B* coding sequence from the WT EMCV clone, with an additional Kozak sequence, the HA tag and a GGSGGS linker sequence as well as a non-viral termination codon between *Pac*I and *Bgl*II restriction sites (sequence available upon request).

For all cellular proteins identified as being potential interaction partners of 2B* (by affinity capture coupled to quantitative proteomics), corresponding coding sequences were cloned from cDNA of uninfected BHK-21 cells. RNA was extracted from BHK-21 cells, reverse transcribed (see above) and the cDNA used for PCR amplification.

Primers for the PCR amplification of all constructs were designed to add to each gene flanking *Pac*I and either *Afl*II or *Bgl*II restriction sites, as well as an N-terminal FLAG tag separated by a GGSGGS linker. These amplicons were then digested and ligated into pCAGG.

### DNA transfection

BHK-21 cells were transfected at 60–70% confluency, in a 6 well plate seeded the day prior. 15 µL lipofectamine 2000 (Invitrogen) was added to 300 µL Opti-MEM and a total of 4 µg DNA was added into another 300 µL Opti-MEM. These solutions were then mixed and incubated for 20 min while cells were washed with PBS. Following addition of transfection mixture, cells were incubated for 3.5 h at 37°C with gentle agitation. 2 mL of complete DMEM was then added to each well and incubated for 24 h. For all co-immunoprecipitation samples, including those used for TMT mass spectrometry analysis, 10 cm dishes of BHK-21 were transfected using a scaled protocol, similar to that described above.

### Frameshifting assays

For the WT sequence and each mutant to be tested, 105 nt (106 nt for −1 SS-SL) covering the 2B* frameshift site (11 nt upstream, 7 nt slippery sequence, 87 nt downstream) was amplified by 2-step overlap extension PCR (primer sequences available upon request, numbers exclude restriction enzyme sites). Amplicons were ligated into pSGDLuc via *Bgl*II and *Psp*XI restriction sites. These fragments included the desired mutations for testing, as well as the slippery sequence and stem-loop required for frameshifting(3).

BHK-21 cells were subcultured 24 h prior to being reverse transfected with the pSGDLuc plasmids in a 96 well plate. Upon reverse transfection, cells were trypsinised and washed in DMEM containing 2% FCS, 25 mM HEPES and 1 mM L-glutamine and washed again in DMEM with 25 mM HEPES and 1 mM L-glutamine without FCS. The cells were then resuspended in this sera-free DMEM. 150 ng of total DNA and 0.5 µL lipofectamine 2000 per well was incubated at room temperature in 10.5 µL of Opti-MEM (Gibco) for 20 min prior to the addition of 6.5 × 10^4^ BHK-21 cells per well. 5% FCS was added, and the transfections were immediately seeded into plates in triplicate and incubated for 24 h, before being frozen in 50 µL of passive lysis buffer (Promega). After thawing, 30 µL of the lysate was sequentially mixed with 30 µL of each luciferase reagent (Promega) as described by the manufacturer.

### Quantitative RT-PCR

Following transfection of MEF cells, the media was changed prior to addition of Opti-MEM with lipofectamine 2000 (Invitrogen) and poly(I:C) to a final concentration of 10 µg/mL poly(I:C). After 6 h RNA was extracted and1 µg of total RNA was used for reverse transcription (Quantitect reverse transcription kit, Qiagen) as per instructions. cDNA was amplified using a ViiA 7 real-time PCR system (ThermoFisher Scientific). The reaction cycle threshold was determined using the following program: i) initial heating to 55°C for 2 min; ii) initial denaturation for 10 min at 95°C; iii) 40 cycles of denaturation for 15 s at 95°C, annealing for 1 min at 60°C. The melt curve was calculated by denaturation for 15 s at 95°C, annealing for 1 min at 60°C with 0.05°C increments to 95°C. The primer sequences for mouse IL6 were (sense) 5’ GAAGTTCCTCTCTGCAAGAGACTTCCATC and (antisense) 5’ GAAGTTCCTCTCTGCAAGAGACTTCCATC. Primer sequences for other transcripts are available upon request and have been previously published (44–46). Primer efficiency was calculated during qRT-PCR optimisation and an efficiency between 90– 110% was obtained for all primer pairs (data not shown). During data analysis, amplification of each gene was normalised to GAPDH amplification. Fold change of each gene was calculated relative to the empty vector transfected, poly(I:C) stimulated control.

### EMCV reverse genetics

The parental (WT) EMCV sequence has been previously described (3) and is based on the EMCV subtype mengovirus cDNA, pMC0, developed by Ann Palmenberg (University of Wisconsin-Madison) (47). It resembles GenBank accession DQ294633.1, although the poly(C) tract is absent and there are 13 single-nucleotide differences (A2669C, G3044C, C3371T, A4910C, G4991A, C5156T, G5289A, G5314C, G5315A, A5844C, G6266A, G6990A, A6992G; DQ294633.1 coordinates). The 2B*KO mutation in a subclone has been previously described (11). Regions encompassing the mutations of interest were digested using *Bgl*II and *Pac*I and ligated into the WT EMCV molecular clone.

The GFP-Lzn EMCV clone was a kind gift from Prof. Frank van Kuppeveld and was derived from the clone pRLuc-QG-M16.1 (13, 48). The Lzn mutation was mutated to WT and the 2B*KO and GNN mutations were subsequently introduced (primer sequences available upon request). All GFP viral clones contain the EMCV IRES followed by the first 6 codons of the viral protein L, followed by sequence encoding GFP inserted into restriction enzyme cloning sites. The viral 3C protease cleavage site Gln-Gly immediately precedes the full-length viral polyprotein coding sequence, including L but without the initiation methionine, allowing the GFP to be separated from L. Sanger sequencing was used to sequence the full length of all mutant and WT viral clones (Biochemistry Sequencing Facility, University of Cambridge).

EMCV molecular clones were linearised with *BamH*I and genomic RNA was *in vitro* transcribed using the T7 RiboMax kit (Promega). Transcripts were purified (RNeasy kit, Qiagen) before being transfected into BHK-21 cells to generate virus stocks.

### Preparation of virus stocks

BHK-21 cells were grown to 60–70% confluency in 10 cm dishes. All Opti-MEM (Gibco) was supplemented with 1:1000 RNaseOUT (Invitrogen). 90 µL lipofectamine 2000 (Invitrogen) was added to 1.8 mL Opti-MEM and 14 µg RNA was added to another 1.8 mL Opti-MEM. These solutions were mixed together and incubated for 20 min while cells were washed with PBS. Following addition of transfection mixture, cells were incubated for 3.5 h at room temperature with gentle agitation. Transfection medium was removed prior to the addition of 8 mL of 2% FCS DMEM and incubation for 24 h or until full CPE was visible. The plates were then scraped and samples freeze/thawed thrice, prior to the dead cells and debris being removed by light clarification and the supernatant frozen at −70°C in aliquots. Virus concentration was estimated by plaque assay.

### Plaque assays

BSR cells were seeded at 35% confluency in 6 well plates 24 h prior to infection. The media was removed and the cells were washed once with PBS before the addition of 1 mL sera-free DMEM containing the specified dilution of virus. The infection was incubated at room temperature with continuous rocking for 1 h before the inocula was removed, the cells were washed once with PBS and 3 mL of DMEM (2% FCS) containing 1% low melting point agarose (ThermoFisher Scientific) was added to each well. Cells were incubated for 48 h before being fixed with 4% formaldehyde and stained with toluidine blue. Plaques were counted manually and their sizes quantified using ImageJ (49, 50).

### Immunoblots

During validation of 2B*KO EMCV, all samples to be analysed by immunoblot were frozen in RIPA buffer (ThermoFisher Scientific) containing 1:10,000 Benzonase nuclease (Sigma) and protease and phosphatase inhibitors (Halt, ThermoFisher Scientific) (“complete RIPA”). All samples were boiled in SDS-based protein loading dye (Laemmli buffer) containing 10 mM DTT for 7 min prior to electrophoresis. Following resolution, proteins were transferred to 0.2 µm nitrocellulose membranes by semi-dry electrotransfer in a Transblot turbo transfer system using recommended standard settings (Bio-Rad). Membranes were blocked with 5% w/v non-fat milk powder in PBS for a minimum of 3 h with continuous rocking. Primary antibodies were diluted in blocking buffer prior to incubation with the membrane for at least 2 h. Membranes were washed 3 times with TBS-Tween20 (0.1%) before being incubated with the relevant secondary antibody for 1 h with continuous rocking. Membranes were again washed 3 times with TBS-Tween20 (0.1%) before being imaged with the Odyssey CLx imaging system (LI-COR).

### Quantitative proteomics and TMT labelling of co-immunoprecipitation samples from infected BHK-21 cells

BHK-21 cells were mock infected or infected with HA tagged WT and HA tagged 2B*KO EMCV at an MOI 5.0 for 10 hpi in five, 10cm dishes per sample, each in triplicate. Due to the number of dishes, samples were separated into two groups and infected one hour apart to ensure the number of IP samples would be manageable. The HA tagged 2B* was then purified using the Pierce Magnetic HA-Tag IP/Co-IP Kit with wash buffer supplemented with Halt protease and phosphatase inhibitors (ThermoFisher Scientific). During the final wash, 12.5% of each sample was retained in separate tubes for western blot analysis. Following the removal of the buffer for the third and final wash from the rest of the sample, the beads were snap frozen. Samples were eluted from the beads by boiling in 2x SDS Loading buffer and prepared for downstream proteomic analysis by SP3 sample preparation (51). In brief, samples were reduced and alkylated, followed by precipitation onto magnetic beads by the addition of ethanol to 80% (v/v) final concentration. Samples were then washed three times in 90% (v/v) ethanol, and then resuspended in 100mM TEAB containing 20ng/uL Trypsin Gold and digested overnight. Following digestion, samples were removed from the beads and labelled with Tandem Mass Tag reagents (ThermoFisher Scientific), before quenching with the addition of hydroxylamine. Samples were desalted by stage-tipping before analysis by LC-MS/MS on a Dionex 3000 coupled in line to a Q-Exactive-HF mass spectrometer using data-dependent acquisition. Analysis was performed using MaxQuant (52) using a fasta file containing Syrian Golden Hamster and EMCV protein sequences. Raw data and the.fasta files used have been uploaded to the PRIDE repository [accession number pending] (53).

Downstream data analysis of the MaxQuant proteinGroups output file was conducted using Perseus 2.0.11 (54). Reverse database hits, common contaminants (MaxQuant contaminant list) and proteins only identified by site were removed. Data were normalised for equal protein loading based on the median total reporter intensity and then subjected to a log_2_(*x*) transformation. Replicate samples were grouped and rows with < 3 valid values in at least one group were removed. Missing data were imputed from the normal distribution before Student’s *t*-tests. Putative interaction partners were defined as having both −log_10_(*p*-value) greater than 1.3 (*p*-value >0.05) and fold change of greater than 2 (log_2_(Fold change > 1) in both the comparison of HA2B* EMCV *vs* mock and HA2B* EMCV *vs* HA2B*KO EMCV.

### Anti-FLAG immunoprecipitation

BHK-21 cells were co-transfected with both pCAGG-HA2B* and a cloned FLAG-tagged putative interaction partner (pCAGG-X) in 10 cm dishes. 24 h after transfection, the cells were washed in ice cold PBS before being scraped and lysed in 750 µL of lysis buffer (25 mM Tris HCl [pH 7.4], 150 mM NaCl, 1 mM EDTA, 1% NP40, 5% glycerol, pH 7.4) for 10 min at 4°C. The lysates were clarified and 50 µL was retained (input). The FLAG-tagged bait protein and any interacting proteins were purified using anti-FLAG resin (Sigma). Briefly, lysates were incubated with the resin with continuous rotation for a minimum of 2 h at 4°C, then washed thrice in fresh lysis buffer. Following the removal of the final wash, the resin was boiled for 5 min in 60 µL 2x protein loading dye (Laemmli buffer) in the absence of DTT. The undissolved resin was removed by centrifugation and DTT was added (final concentration 5 mM) prior to reheating the sample for a further 3 min. Proteins were resolved by polyacrylamide gel electrophoresis and imaged by immunoblot against the epitope tags.

### Confocal microscopy

BSR cells were seeded at 20% confluency on to glass coverslips in 24 well plates, 24 h prior to transfection (see above). 24 h after transfection, cells were washed in PBS and fixed with 4% paraformaldehyde for 20 min at room temperature. Cells were washed thrice both prior to and immediately following permeabilisation with 0.5% Triton X-100 for 10 min. To reduce nonspecific binding, cells were blocked with 10% BSA for a minimum of 2 h at room temperature with gentle agitation before addition of the relevant primary antibodies. Antibodies against the HA epitope (C29F4, Cell Signaling Technologies) and the FLAG epitope (F1804, Sigma) were each diluted 1:1000 and the coverslips were incubated at 4°C overnight with continuous gentle rocking. Samples were thrice washed in PBS before the addition of fluorophore-labelled secondary antibodies diluted in 10% BSA (anti-rabbit Alexa-fluor 488 (Invitrogen); anti-rat Alexa-fluor 594 (abcam)). Samples were again incubated with continuous rocking at room temperature for 1 h. Finally, samples were thrice washed and the coverslips were mounted upon glass slides. DAPI was included in the mounting media (Prolong gold antifade with DAPI, ThermoFisher Scientific). Samples were imaged at 63 x on a LSM 700 laser scanning confocal microscope.

### Live cell imaging

BHK-21 cells infected with GFP-tagged viruses were imaged at regular intervals using the Incucyte live cell analysis system and the Incucyte 2022B Rev2 GUI software (Sartorius). Three regions were imaged per well using a 10x objective (96 well plate, each sample in technical triplicate). The background threshold for each plate was independently set using manually selected images, with high and low fluorescence levels. Cell confluence was measured by phase microscopy. GCU at each time point was normalised to the respective GCU in that well at 5.5 h post infection.

### *In silico* protein modelling

Alphafold (55), I-TASSER (56) and Netsurfp 3.0 (57) were used to predict the secondary structure of 2B*, using the amino acid sequence from GenBank accession DQ294633.1.

## ACKNOWLEDGEMENTS

The authors would like to thank Prof. Frank J.M. van Kuppeveld (Utrecht University) for kindly providing a molecular clone of GFP-tagged EMCV. The authors would like to thank Prof. Ian Brierley, Prof. Colin Crump and Prof. Stephen Graham (University of Cambridge) for useful discussions, as well as Prof. Ann Palmenberg (University of Wisconsin-Madison) for providing additional supporting reagents. A.E.F. and H.S. were supported by Wellcome Trust Senior Research Fellowships (106207/Z/14/Z, 220814/Z/20/Z) and a European Research Council grant (646891). J.R.E. was supported by a Sir Henry Dale Fellowship jointly funded by the Wellcome Trust and the Royal Society (216370/Z/19/Z). EE and SH are supported by funding from the Academy of Medical Sciences (SBF006\1008), the Medical Research Council (MR/X000885/1), and a Wellcome Trust Career Development Award (227831/Z/23/Z) awarded to EE. S.K.N. was supported by a Department of Pathology, University of Cambridge PhD studentship that was funded by the Waldmann Fund. The funders had no role in study design or data interpretation, or the decision to submit the work for publication. For the purposes of open access, the authors have applied a CC BY public copyright licence to any Author Accepted article version arising from this submission. Figure 6 was created with BioRender.com and is published under Agreement Number ZR26VGVQ7I.

## Author Contributions

S.K.N., H.S. and A.E.F. designed the study and experiments. S.K.N., E.E., N.L., L.G.C. and S.H. performed experiments. H.S., J.R.E., H.B., I.G., A.S.J., E.E. and A.E.F. provided expertise on data analysis, interpretation and methodology. S.K.N. and H.S. wrote the article. All authors edited and discussed the article. A.E.F., J.R.E. and E.E. secured funding.

## TABLE TITLES AND LEGENDS

**Table 1:** Proteins which were statistically significantly enriched in HA2B* EMCV infected samples relative to both HA2B*KO EMCV infected and mock infected samples.

## SUPPLEMENTAL MATERIAL

**Figure S1:** BSR cells were co-transfected with pCAGG-HA2B* and either the gene encoding 14-3-3β or 14-3-3γ, with an N-terminal FLAG tag, also in a pCAGG expression construct. Cells were fixed at 24 h and stained with both anti-HA and anti-FLAG antibodies. Nuclei were stained with DAPI during mounting. Images are representative of three independent biological repeats.

**Figure S2:** BSR cells were co-transfected with pCAGG-HA2B* and the gene encoding 14-3-3ε (isoform X1), 14-3-3ζ or 14-3-3η with an N-terminal FLAG tag, also in a pCAGG expression construct. Cells were fixed at 24 h and stained with both anti-HA and anti-FLAG antibodies. Nuclei were stained with DAPI during mounting. Images are representative of three independent biological repeats.

**Figure S3:** BSR cells were co-transfected with pCAGG-HA2B* and the gene encoding 14-3-3η, 14-3-3θ or 14-3-3σ, with an N-terminal FLAG tag, also in a pCAGG expression construct. Cells were fixed at 24 h and stained with both anti-HA and anti-FLAG antibodies. Nuclei were stained with DAPI during mounting. Images are representative of three independent biological repeats.

## REFERENCES

1. Loughran G, Libbey JE, Uddowla S, Scallan MF, Ryan MD, Fujinami RS, Rieder E, Atkins JF. 2013. Theiler’s murine encephalomyelitis virus contrasts with encephalomyocarditis and foot-and-mouth disease viruses in its functional utilization of the StopGo non-standard translation mechanism. J Gen Virol 94:348–353.

2. Loughran G, Firth AE, Atkins JF. 2011. Ribosomal frameshifting into an overlapping gene in the 2B-encoding region of the cardiovirus genome. Proc Natl Acad Sci U S A 108:E1111–9.

3. Napthine S, Ling R, Finch LK, Jones JD, Bell S, Brierley I, Firth AE. 2017. Protein-directed ribosomal frameshifting temporally regulates gene expression. Nat Commun 8:15582.

4. Hill CH, Pekarek L, Napthine S, Kibe A, Firth AE, Graham SC, Caliskan N, Brierley I. 2021. Structural and molecular basis for Cardiovirus 2A protein as a viral gene expression switch. Nat Commun 12:7166.

5. Finch LK, Ling R, Napthine S, Olspert A, Michiels T, Lardinois C, Bell S, Loughran G, Brierley I, Firth AE. 2015. Characterization of Ribosomal Frameshifting in Theiler’s Murine Encephalomyelitis Virus. J Virol 89:8580–9.

6. Hill CH, Cook GM, Napthine S, Kibe A, Brown K, Caliskan N, Firth AE, Graham SC, Brierley I. 2021. Investigating molecular mechanisms of 2A-stimulated ribosomal pausing and frameshifting in Theilovirus. Nucleic Acids Res 49:11938–11958.

7. Tam EH, Liu YC, Woung CH, Liu HM, Wu GH, Wu CC, Kuo RL. 2021. Role of the Chaperone Protein 14-3-3epsilon in the Regulation of Influenza A Virus-Activated Beta Interferon. J Virol 95:e0023121.

8. Riedl W, Acharya D, Lee JH, Liu G, Serman T, Chiang C, Chan YK, Diamond MS, Gack MU. 2019. Zika Virus NS3 Mimics a Cellular 14-3-3-Binding Motif to Antagonize RIG-I-and MDA5-Mediated Innate Immunity. Cell Host Microbe 26:493–503 e6.

9. Gupta S, Yla-Anttila P, Sandalova T, Sun R, Achour A, Masucci MG. 2019. 14-3-3 scaffold proteins mediate the inactivation of trim25 and inhibition of the type I interferon response by herpesvirus deconjugases. PLoS Pathog 15:e1008146.

10. Andrews DDT, Vlok M, Akbari Bani D, Hay BN, Mohamud Y, Foster LJ, Luo H, Overall CM, Jan E. 2023. Cleavage of 14-3-3epsilon by the enteroviral 3C protease dampens RIG-I-mediated antiviral signaling. J Virol 97:e0060423.

11. Ling R, Firth AE. 2017. An analysis by metabolic labelling of the encephalomyocarditis virus ribosomal frameshifting efficiency and stimulators. J Gen Virol 98:2100–2105.

12. Loughran G, Howard MT, Firth AE, Atkins JF. 2017. Avoidance of reporter assay distortions from fused dual reporters. RNA 23:1285–1289.

13. Dorobantu CM, Albulescu L, Harak C, Feng Q, van Kampen M, Strating JR, Gorbalenya AE, Lohmann V, van der Schaar HM, van Kuppeveld FJ. 2015. Modulation of the Host Lipid Landscape to Promote RNA Virus Replication: The Picornavirus Encephalomyocarditis Virus Converges on the Pathway Used by Hepatitis C Virus. PLoS Pathog 11:e1005185.

14. Pennington KL, Chan TY, Torres MP, Andersen JL. 2018. The dynamic and stress-adaptive signaling hub of 14-3-3: emerging mechanisms of regulation and context-dependent protein-protein interactions. Oncogene 37:5587–5604.

15. Liou JY, Ghelani D, Yeh S, Wu KK. 2007. Nonsteroidal anti-inflammatory drugs induce colorectal cancer cell apoptosis by suppressing 14-3-3epsilon. Cancer Res 67:3185–91.

16. Hermeking H, Benzinger A. 2006. 14-3-3 proteins in cell cycle regulation. Semin Cancer Biol 16:183–92.

17. Zhao S, Li B, Li C, Gao H, Miao Y, He Y, Wang H, Gong L, Li D, Zhang Y, Feng J. 2019. The Apoptosis Regulator 14-3-3eta and Its Potential as a Therapeutic Target in Pituitary Oncocytoma. Front Endocrinol (Lausanne) 10:797.

18. Eckardt NA. 2001. Transcription factors dial 14-3-3 for nuclear shuttle. Plant Cell 13:2385–9.

19. Lin JP, Fan YK, Liu HM. 2019. The 14-3-3eta chaperone protein promotes antiviral innate immunity via facilitating MDA5 oligomerization and intracellular redistribution. PLoS Pathog 15:e1007582.

20. Liu HM, Loo YM, Horner SM, Zornetzer GA, Katze MG, Gale M, Jr. 2012. The mitochondrial targeting chaperone 14-3-3epsilon regulates a RIG-I translocon that mediates membrane association and innate antiviral immunity. Cell Host Microbe 11:528–37.

21. Jia H, Liang Z, Zhang X, Wang J, Xu W, Qian H. 2017. 14-3-3 proteins: an important regulator of autophagy in diseases. Am J Transl Res 9:4738–4746.

22. Gu Y, Xu K, Torre C, Samur M, Barwick BG, Rupji M, Arora J, Neri P, Kaufman J, Nooka A, Bernal-Mizrachi L, Vertino P, Sun SY, Chen J, Munshi N, Fu H, Kowalski J, Boise LH, Lonial S. 2018. 14-3-3zeta binds the proteasome, limits proteolytic function and enhances sensitivity to proteasome inhibitors. Leukemia 32:744–751.

23. Kumar M, Michael S, Alvarado-Valverde J, Meszaros B, Samano-Sanchez H, Zeke A, Dobson L, Lazar T, Ord M, Nagpal A, Farahi N, Kaser M, Kraleti R, Davey NE, Pancsa R, Chemes LB, Gibson TJ. 2022. The Eukaryotic Linear Motif resource: 2022 release. Nucleic Acids Res 50:D497–D508.

24. Coblitz B, Wu M, Shikano S, Li M. 2006. C-terminal binding: an expanded repertoire and function of 14-3-3 proteins. FEBS Lett 580:1531–5.

25. Muslin AJ, Tanner JW, Allen PM, Shaw AS. 1996. Interaction of 14-3-3 with signaling proteins is mediated by the recognition of phosphoserine. Cell 84:889–97.

26. Gack MU, Shin YC, Joo CH, Urano T, Liang C, Sun L, Takeuchi O, Akira S, Chen Z, Inoue S, Jung JU. 2007. TRIM25 RING-finger E3 ubiquitin ligase is essential for RIG-I-mediated antiviral activity. Nature 446:916–920.

27. Reikine S, Nguyen JB, Modis Y. 2014. Pattern Recognition and Signaling Mechanisms of RIG-I and MDA5. Front Immunol 5:342.

28. Seth RB, Sun L, Ea CK, Chen ZJ. 2005. Identification and characterization of MAVS, a mitochondrial antiviral signaling protein that activates NF-kappaB and IRF 3. Cell 122:669–82.

29. Li L, Fan H, Song Z, Liu X, Bai J, Jiang P. 2019. Encephalomyocarditis virus 2C protein antagonizes interferon-beta signaling pathway through interaction with MDA5. Antiviral Res 161:70–84.

30. Hato SV, Ricour C, Schulte BM, Lanke KH, de Bruijni M, Zoll J, Melchers WJ, Michiels T, van Kuppeveld FJ. 2007. The mengovirus leader protein blocks interferon-alpha/beta gene transcription and inhibits activation of interferon regulatory factor 3. Cell Microbiol 9:2921–30.

31. Stavrou S, Feng Z, Lemon SM, Roos RP. 2010. Different strains of Theiler’s murine encephalomyelitis virus antagonize different sites in the type I interferon pathway. J Virol 84:9181–9.

32. Manivannan P, Siddiqui MA, Malathi K. 2020. RNase L Amplifies Interferon Signaling by Inducing Protein Kinase R-Mediated Antiviral Stress Granules. J Virol 94.

33. Drappier M, Jha BK, Stone S, Elliott R, Zhang R, Vertommen D, Weiss SR, Silverman RH, Michiels T. 2018. A novel mechanism of RNase L inhibition: Theiler’s virus L* protein prevents 2-5A from binding to RNase L. PLoS Pathog 14:e1006989.

34. Brunet A, Kanai F, Stehn J, Xu J, Sarbassova D, Frangioni JV, Dalal SN, DeCaprio JA, Greenberg ME, Yaffe MB. 2002. 14-3-3 transits to the nucleus and participates in dynamic nucleocytoplasmic transport. J Cell Biol 156:817–28.

35. Jen G, Thach RE. 1982. Inhibition of host translation in encephalomyocarditis virus-infected L cells: a novel mechanism. J Virol 43:250–61.

36. Bustos DM. 2012. The role of protein disorder in the 14-3-3 interaction network. Mol Biosyst 8:178–84.

37. Rogers KJ, Jones-Burrage S, Maury W, Mukhopadhyay S. 2020. TF protein of Sindbis virus antagonizes host type I interferon responses in a palmitoylation-dependent manner. Virology 542:63–70.

38. Khalil AM, Nogales A, Martinez-Sobrido L, Mostafa A. 2024. Antiviral responses versus virus-induced cellular shutoff: a game of thrones between influenza A virus NS1 and SARS-CoV-2 Nsp1. Front Cell Infect Microbiol 14:1357866.

39. Zhang B, Liu M, Huang J, Zeng Q, Zhu Q, Xu S, Chen H. 2022. H1N1 Influenza A Virus Protein NS2 Inhibits Innate Immune Response by Targeting IRF7. Viruses 14.

40. da Costa LS, Outlioua A, Anginot A, Akarid K, Arnoult D. 2019. RNA viruses promote activation of the NLRP3 inflammasome through cytopathogenic effect-induced potassium efflux. Cell Death Dis 10:346.

41. Schwarz EM, Badorff C, Hiura TS, Wessely R, Badorff A, Verma IM, Knowlton KU. 1998. NF-kappaB-mediated inhibition of apoptosis is required for encephalomyocarditis virus virulence: a mechanism of resistance in p50 knockout mice. J Virol 72:5654–60.

42. Li Y, Shang P, Shyu D, Carrillo C, Naraghi-Arani P, Jaing CJ, Renukaradhya GJ, Firth AE, Snijder EJ, Fang Y. 2018. Nonstructural proteins nsp2TF and nsp2N of porcine reproductive and respiratory syndrome virus (PRRSV) play important roles in suppressing host innate immune responses. Virology 517:164–176.

43. Pavesi A. 2021. Origin, Evolution and Stability of Overlapping Genes in Viruses: A Systematic Review. Genes (Basel) 12.

44. Jahun AS, Sorgeloos F, Chaudhry Y, Arthur SE, Hosmillo M, Georgana I, Izuagbe R, Goodfellow IG. 2023. Leaked genomic and mitochondrial DNA contribute to the host response to noroviruses in a STING-dependent manner. Cell Rep 42:112179.

45. Petro TM. 2005. ERK-MAP-kinases differentially regulate expression of IL-23 p19 compared with p40 and IFN-beta in Theiler’s virus-infected RAW264.7 cells. Immunol Lett 97:47–53.

46. Wu Y, Li YY, Matsushima K, Baba T, Mukaida N. 2008. CCL3-CCR5 axis regulates intratumoral accumulation of leukocytes and fibroblasts and promotes angiogenesis in murine lung metastasis process. J Immunol 181:6384–93.

47. Duke GM, Osorio JE, Palmenberg AC. 1990. Attenuation of Mengo virus through genetic engineering of the 5’ noncoding poly(C) tract. Nature 343:474–6.

48. Albulescu L, Wubbolts R, van Kuppeveld FJ, Strating JR. 2015. Cholesterol shuttling is important for RNA replication of coxsackievirus B3 and encephalomyocarditis virus. Cell Microbiol 17:1144–56.

49. Schindelin J, Arganda-Carreras I, Frise E, Kaynig V, Longair M, Pietzsch T, Preibisch S, Rueden C, Saalfeld S, Schmid B, Tinevez JY, White DJ, Hartenstein V, Eliceiri K, Tomancak P, Cardona A. 2012. Fiji: an open-source platform for biological-image analysis. Nat Methods 9:676-82.

50. Rueden CT, Schindelin J, Hiner MC, DeZonia BE, Walter AE, Arena ET, Eliceiri KW. 2017. ImageJ2: ImageJ for the next generation of scientific image data. BMC Bioinformatics 18:529.

51. Hughes CS, Moggridge S, Muller T, Sorensen PH, Morin GB, Krijgsveld J. 2019. Single-pot, solid-phase-enhanced sample preparation for proteomics experiments. Nat Protoc 14:68–85.

52. Tyanova S, Temu T, Cox J. 2016. The MaxQuant computational platform for mass spectrometry-based shotgun proteomics. Nat Protoc 11:2301–2319.

53. Perez-Riverol Y, Bai J, Bandla C, Garcia-Seisdedos D, Hewapathirana S, Kamatchinathan S, Kundu DJ, Prakash A, Frericks-Zipper A, Eisenacher M, Walzer M, Wang S, Brazma A, Vizcaino JA. 2022. The PRIDE database resources in 2022: a hub for mass spectrometry-based proteomics evidences. Nucleic Acids Res 50:D543–D552.

54. Tyanova S, Temu T, Sinitcyn P, Carlson A, Hein MY, Geiger T, Mann M, Cox J. 2016. The Perseus computational platform for comprehensive analysis of (prote)omics data. Nat Methods 13:731–40.

55. Jumper J, Evans R, Pritzel A, Green T, Figurnov M, Ronneberger O, Tunyasuvunakool K, Bates R, Zidek A, Potapenko A, Bridgland A, Meyer C, Kohl SAA, Ballard AJ, Cowie A, Romera-Paredes B, Nikolov S, Jain R, Adler J, Back T, Petersen S, Reiman D, Clancy E, Zielinski M, Steinegger M, Pacholska M, Berghammer T, Bodenstein S, Silver D, Vinyals O, Senior AW, Kavukcuoglu K, Kohli P, Hassabis D. 2021. Highly accurate protein structure prediction with AlphaFold. Nature 596:583–589.

56. Yang J, Yan R, Roy A, Xu D, Poisson J, Zhang Y. 2015. The I-TASSER Suite: protein structure and function prediction. Nat Methods 12:7–8.

57. Hoie MH, Kiehl EN, Petersen B, Nielsen M, Winther O, Nielsen H, Hallgren J, Marcatili P. 2022. NetSurfP-3.0: accurate and fast prediction of protein structural features by protein language models and deep learning. Nucleic Acids Res 50:W510–W515.

