## Supplementary material for "Encephalomyocarditis virus protein 2B* antagonises innate immune signalling by interacting with 14-3-3 protein family members": Table 1

| Protein | $Log_2$(Fold change WT/KO) | $-Log_{10}$(Student's *t*-test *p*-value) WT/KO | $Log_2$(Fold change WT/Mock) | $-Log_{10}$(Student's *t*-test *p*-value) WT/Mock |
|---|---|---|---|---|
| EMCV 3D | 1.01 | 2.21 | 1.33 | 2.37 |
| Safb1 | 1.06 | 3.45 | 1.26 | 4.00 |
| S100A16 | 1.14 | 3.41 | 1.04 | 2.74 |
| S100A13 | 1.27 | 1.70 | 1.06 | 1.93 |
| EMCV 3C | 1.40 | 2.83 | 2.23 | 2.94 |
| Annexin A2 | 1.42 | 3.37 | 1.36 | 2.85 |
| EMCV 2C | 1.65 | 2.39 | 1.65 | 2.39 |
| Calpactin I light chain (S100A10) | 2.41 | 3.41 | 2.71 | 2.61 |
| 14-3-3β | 2.44 | 3.26 | 2.51 | 3.22 |
| EMCV 1B | 2.84 | 3.47 | 5.37 | 4.50 |
| 14-3-3ε | 3.16 | 4.44 | 3.19 | 4.31 |
| 14-3-3γ | 3.26 | 4.44 | 3.37 | 4.24 |
| 14-3-3η | 3.40 | 5.02 | 3.38 | 4.94 |
| 14-3-3ζ | 3.56 | 4.75 | 3.56 | 4.71 |
| 14-3-3θ | 3.78 | 4.20 | 3.86 | 4.14 |
| EMCV 2B* | 3.81 | 4.80 | 3.85 | 4.81 |
| 14-3-3σ | 3.98 | 3.91 | 4.21 | 4.96 |
