## Supplementary figures and images for "Encephalomyocarditis virus protein 2B* antagonises innate immune signalling by interacting with 14-3-3 protein family members"

### Supplemental Figure 1

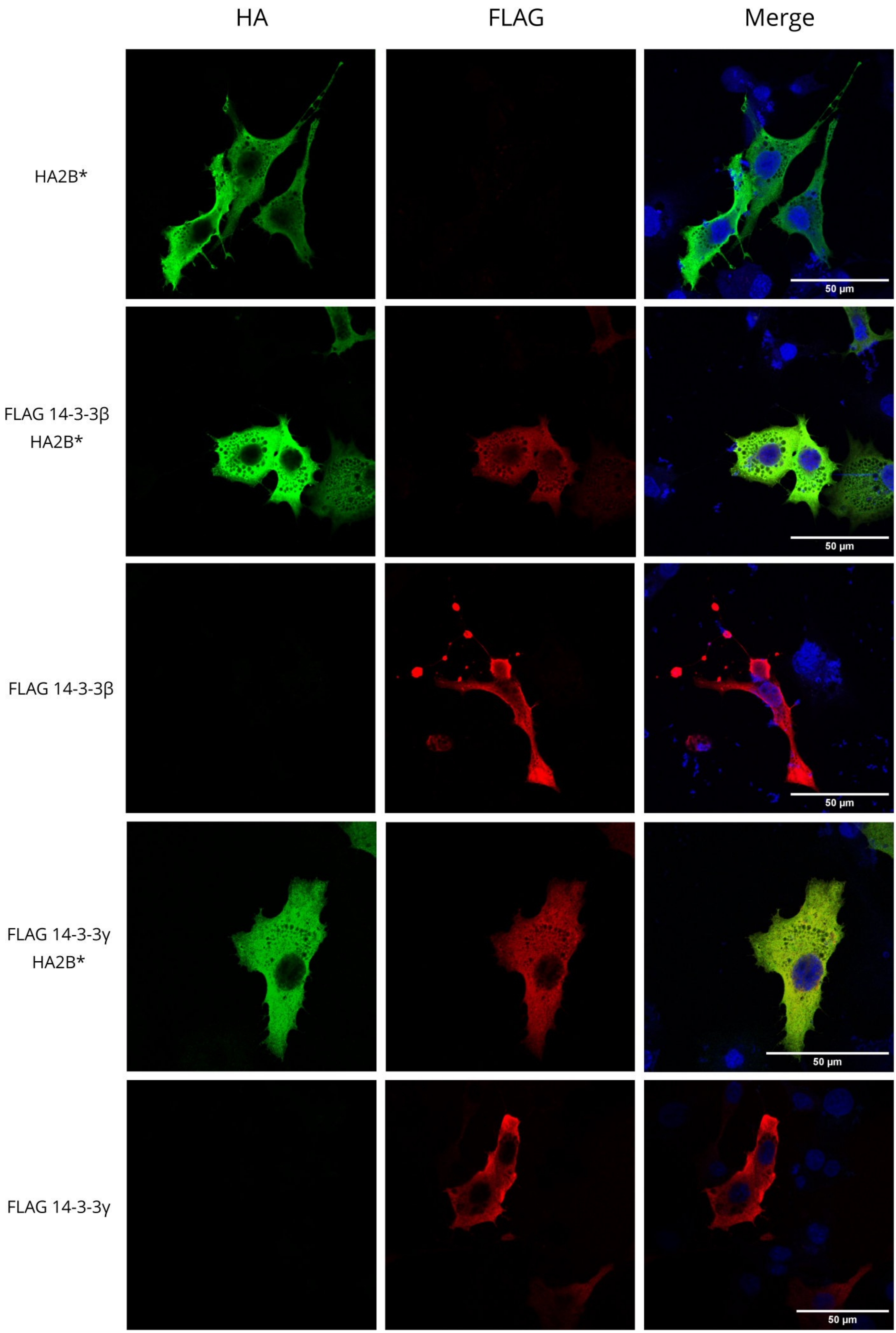

### Supplemental Figure 2

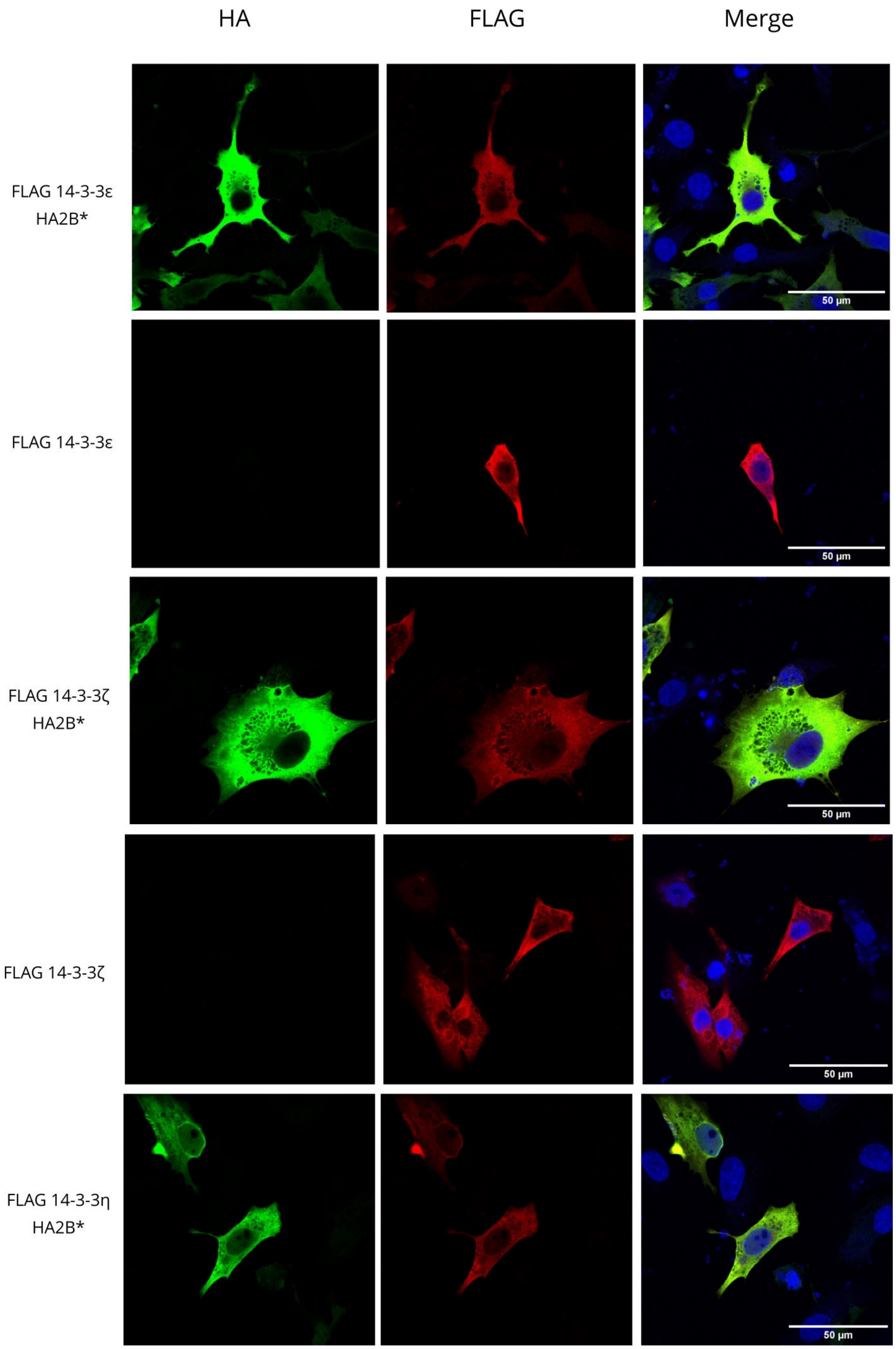

### Supplemental Figure 3

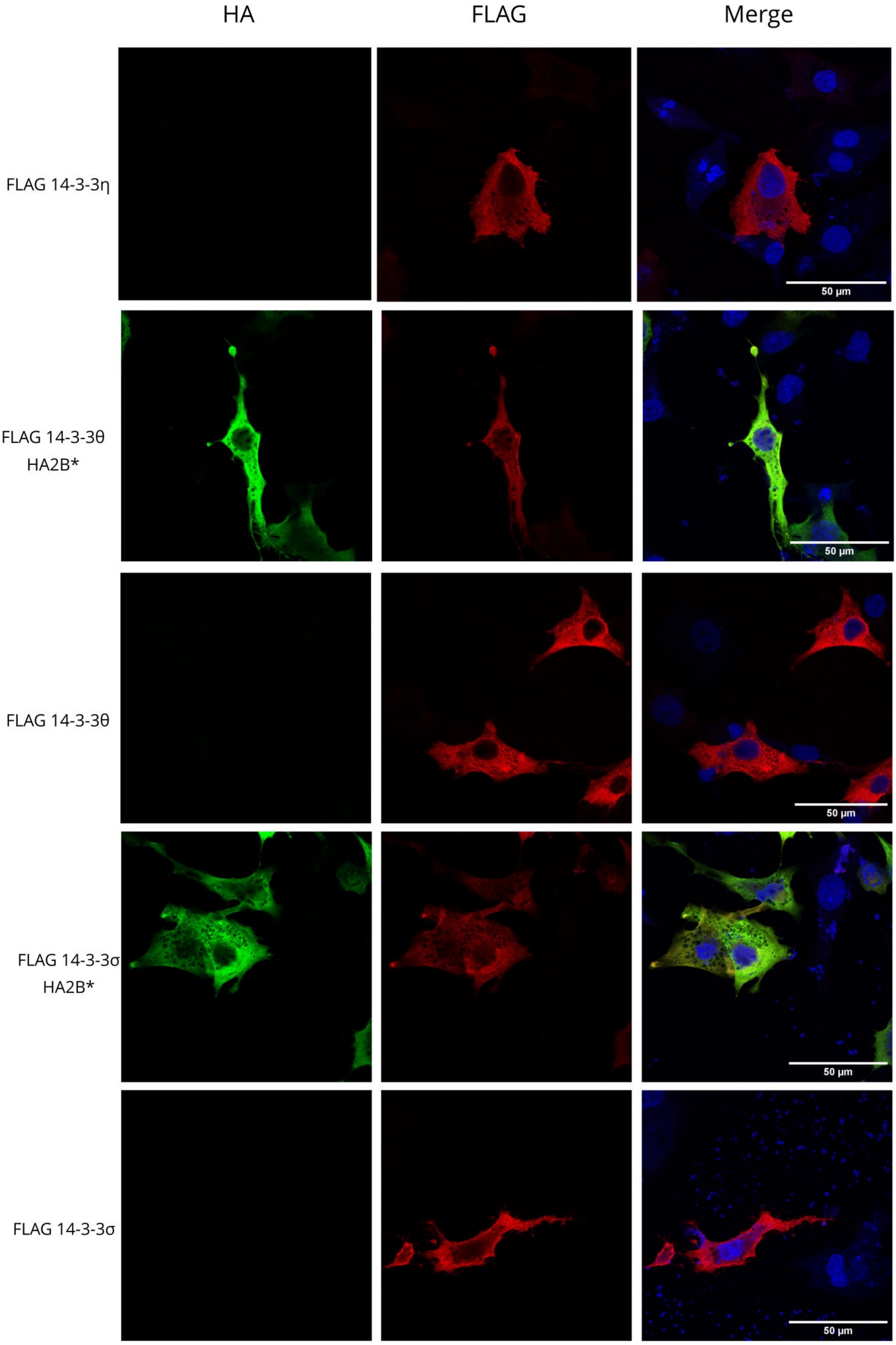
